# Topoisomerase IIα orchestrates secretion of IL-6 and IL-8 with human papillomavirus replication

**DOI:** 10.1101/2025.04.03.647014

**Authors:** Yanfei Liu, Zi Han, Paul Kaminski, Chengcheng Tao, Xiaoge Li, Mengmeng Liu, Yang Li, Ying Jia, Junfen Xu, Shiyuan Hong

## Abstract

High-risk human papillomavirus (HPV) replication requires deregulation of host DNA damage response (DDR) and inflammatory pathways. DNA topoisomerase 2β (Top2β) was previously shown to promote HPV replication, while the function of its paralog Top2α in the viral life cycle remains unknown. Elevated levels of Top2α are consistently observed in cervical intraepithelial lesions and the related carcinomas, as well as in HPV-positive cell lines. Silencing Top2α with shRNA severely suppresses HPV genome maintenance and amplification, but in a DDR-independent manner. Instead, Top2α facilitates secretion of interleukin (IL)-6 and IL-8, which are necessary for HPV replication. Mechanistically, this manipulation is regulated by toll-like receptor 4 (TLR4). Top2α binds to the TLR4 promoter to transcriptionally induce TLR4 expression. Blockade of TLR4 signaling by the specific inhibitor TAK-242 significantly reduces the secreted IL-6/IL-8 levels and HPV replication. Overall, our results reveal a novel role of Top2α to shape the inflammatory microenvironment that benefits HPV replication, making it a promising therapeutic target for HPV-associated diseases.

**IMPORTANCE:** Human papillomaviruses (HPVs) are oncogenic pathogens responsible for many HPV-related diseases such as cervical cancer. HPV replication depends on the subversion of host cellular machineries, particularly pathways governing DNA damage response (DDR) and inflammation. However, the roles of enzymes that directly introduce DNA nicks or mediate strand rejoining remain poorly characterized. Here we highlight a novel role of DNA topoisomerase 2α (Top2α) in promoting HPV genome maintenance and amplification but in a DDR-independent manner. Unlike its previously studied paralog Top2β, Top2α supports HPV replication by driving the secretion of inflammatory molecules interleukin (IL)-6 and IL-8, which are critical for the viral genome persistence and copy number expansion. To our knowledge, this is the first evidence that a DDR-associated protein manipulates cytokine secretion, rather than expression, to support HPV replication. Furthermore, blocking toll-like receptor 4 (TLR4) activities significantly reduced Top2α-dependent IL-6/IL-8 secretion and impaired HPV replication. These findings establish Top2α as a crucial player in HPV pathogenesis, with a therapeutic potential for treating HPV-associated malignancies.

## INTRODUCTION

Human papillomaviruses (HPVs) are non-enveloped double-stranded, circular DNA viruses, associated with approximately 5% of human cancers (1). The productive replication of HPVs is intricately dependent on differentiation of the infected epithelial cells (2, 3), and requires activation of host DNA damage response (DDR) and deregulation of inflammatory response (4–6). DDR is a coordinated signaling network of host defense triggered by DNA damage, characterized by the activation of the key DDR kinases such as the ataxia-telangiectasia-mutated (ATM), ATM and Rad3-related (ATR), and DNA-dependent protein kinase catalytic subunit (DNA-PKcs). ATM or ATR activation has been shown to be essential for HPV replication (7, 8). The associated DDR factors of the ATM/ATR pathways, such as BRD4 (9), 53BP1/BARD1 (10), SIRT1 (11), SMC5/6 (12), RNF168 (13), NBS1 (14), and topoisomerase IIβ-binding protein 1 (TopBP1) (15, 16), also play critical roles in the viral production. Despite these studies, the roles of other DDR regulators in HPV replication still remain to be extensively investigated.

DNA topoisomerase IIα (Top2α) and topoisomerase IIβ (Top2β), which are bound to TopBP1 (17–20), are important enzymes that regulate the DNA topology during replication and transcription by alleviating DNA supercoils or torsional stress at topologically associated domains (21, 22). While Top2β has been shown to be necessary for HPV genome amplification and maintenance (23), the function of its paralog, Top2α, is unknown in the context of HPV replication. Top2α is essential for chromosome condensation, DNA replication, and mitotic chromosome stability (24, 25). Recent studies have shown that Top2α plays a significant role in stalled fork reversal by resolving topological barriers and recruiting DNA translocase PICH (26). Top2α also regulates SMC5/6 complex to promote a G2 arrest (27). Inhibition of Top2α constrains the development of social behavior of zebrafish via PRC2 and H3K27me3 signaling (28). In addition, Top2α has been indicated to be a potential biomarker of cervical premalignant lesions (29), raising the question of whether Top2α plays significant roles in HPV replication or HPV tumorigenesis.

As a hallmark of many viral infections (30, 31), inflammation has been shown to be involved in the regulation of HPV genome maintenance and amplification (32). For example, interferon (IFN)-κ reduces HPV viral transcription via Sp100 restriction capabilities (33). IFN-γ has been demonstrated to suppress HPV amplification in a signal transducer and activator of transcription 1 (STAT1)-dependent manner (34). In contrast, the pro-inflammatory interleukin-8 (IL-8, also referred as CXCL8) is induced by STAT5 activation in HPV-positive cells and promotes HPV replication upon epithelial differentiation (35). IL-6 is capable of activating the STAT3 signaling pathway, which promotes survival and proliferation of cervical cancer cells (36). However, its role in HPV replication is not well understood. Additionally, previous studies demonstrated that disruption of ATR (37) or TopBP1 (15) in HPV-positive cells reprograms expression of inflammatory genes such as IFN-κ and IL-6, possibly via GATA4-dependent or E2F1-dependent manners.

Another critical protein that regulates inflammatory gene expression is toll-like receptor 4 (TLR4), a pattern recognition receptor that acts indispensably in innate immunity and triggers downstream signaling cascades (38). The functions of TLR4 are controversial when hosts were infected with different viruses. The Kasper group showed TLR4-TRIF signaling contributes to gut commensal microbe-induced resistance against vesicular stomatitis virus or influenza (39). Other studies demonstrated TLR4 facilitates replication of influenza A virus (IAV) to induce lung damage (40) or inflammatory response induced by Chikungunya virus (CHIKV) (41). Previous studies have indicated that TLR4 activation leads to the production of pro-inflammatory cytokines such as IL-6 and IL-8 (42), which might be dependent on NF-κB activation (43) or MyD88 and TRIF action (44). Additionally, TLR4 has been shown to promote cervical cancer cell proliferation and apoptosis resistance (45). These studies suggest that TLR4 might contribute to HPV replication by modulating IL-6/IL-8 signaling during HPV life cycle.

In this study, we investigate the role of Top2α in HPV replication, with a particular focus on its involvement in inflammation. Our findings reveal that Top2α is significantly upregulated in cervical cancer tissues and HPV-positive keratinocytes. Silencing Top2α leads to a substantial reduction in the abundance of the viral episomal genomes in both undifferentiating and differentiating HPV-positive cells, without affecting generation or repair of DNA breaks. RNA-seq analysis identified inflammation-related genes, such as IL-6 and IL-8, as the potential downstream targets of Top2α. Further experiments show that Top2α regulates the secretion, but not the expression, of these cytokines in HPV-positive cells through TLR4 signaling. Inhibition of TLR4 expression effectively suppresses HPV replication as well as secretion of IL-6/IL-8 in a dose-dependent manner. Finally, exogenous addition of IL-6 or IL-8 promotes HPV genome maintenance or amplification. These results unveil a previously unrecognized role of Top2α in HPV replication, which extends its canonical functions in DNA topology to the regulation of inflammatory cytokine secretion that benefits viral replication. Our study provides new insights into the interplay between DDR and inflammation during HPV life cycle, suggesting that targeting this crosstalk could offer novel therapeutic strategies against HPV-related cancers.

## RESULTS

### Top2α is upregulated in cervical cancer tissues and HPV-positive cells

We first examined the expression of Top2α in HPV-driven cervical diseases. The Gene expression profiling interactive analysis (GEPIA) database predicted that the TOP2A mRNA levels in cervical cancer tissues are significantly higher than in normal tissues (Fig. 1A). This prediction was validated using RT-qPCR on the tissue samples collected from cervical cancer patients (Fig. 1B). Furthermore, immunohistochemistry (IHC) analysis illustrated that Top2α expression was gradually increased in low-grade squamous intraepithelial lesions (LSIL), high-grade squamous intraepithelial lesions (HSIL), and cervical cancer tissues (Fig. 1C). These observations were then verified in HPV-positive keratinocytes that are derived from human foreskin keratinocytes (HFKs) or the immortalized HaCaT keratinocytes. We found that Top2α levels were greatly elevated in HPV-positive cells that stably harbor either HPV16 (HFK16 and HaCaT16) or HPV31 (HFK31, CIN612, and HaCaT31) episomes, as shown by western blot (Fig. 1D). CIN612 cells were derived from an HPV31-positive biopsy specimen. These data strongly suggest an indispensable role for Top2α in HPV life cycle.

**Fig 1.**
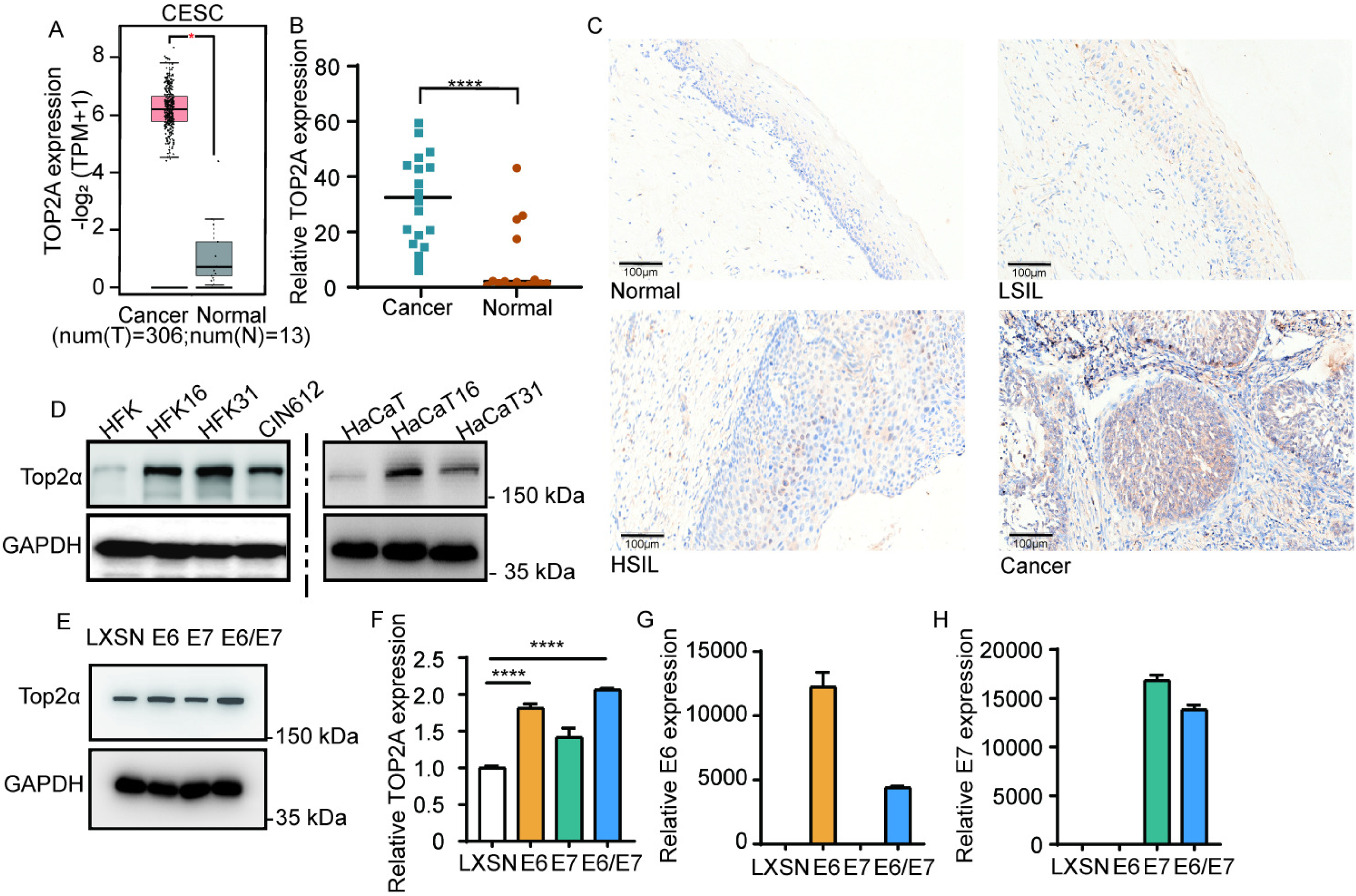
Top2α is upregulated in cervical cancer tissues and HPV-positive cells. (A) The mRNA levels of TOP2A in normal tissues (n=13) and CC tissues (n=306) were compared using the GEPIA database. (B) RT-qPCR analysis was performed to validate the expression of Top2α in 21 normal tissues (Normal) and 20 cervical cancers (Cancer). The relative mRNA levels were calculated using the 2^-ΔΔCT^ method. EEF1A1 was used as the internal control for data normalization. (C) Representative immunohistochemical staining of Top2α in 10 normal tissues, 10 cervical intraepithelial neoplasia, and 10 cervical cancers. (D) Western blot analysis of Top2α levels were characterized in primary human foreskin keratinocytes (HFKs), HFK16 (HPV16 positive), HFK31 (HPV31 positive), and CIN612 (HPV31 positive) (left). Top2α protein levels were tested by western blot in HaCaT, HaCaT16 (HPV16 positive), and HaCaT31 (HPV31 positive) cells (right). GAPDH served as the loading control. Similar results were seen in three independent experiments. The statistical analysis was assayed by 2-tailed t test. \**P*≤0.05, \*\*\*\**P*≤0.0001. (E) Western blot analysis was performed using antibodies of Top2α and GAPDH as the loading control. (F-H) RT-qPCR was performed to detect TOP2A, E6, and E7 mRNA levels in E6, E7, and E6/E7-overexpressing cells.

We have shown that the levels of Top2α are increased in HPV-positive cells (Fig. 1D). Whether this upregulation is due to HPV oncoprotein E6 or E7 remains unclear. To test it, HPV31 E6 or E7 was overexpressed and selected by neomycin resistance in HaCaT cells. Western blot showed that only E6 but not E7 resulted in a modest increase in Top2α expression at protein levels (Fig. 1E). The similar phenomenon was observed by RT-qPCR (Fig. 1F). The expression of E6 and E7 were verified at the mRNA levels by RT-qPCR (Fig. 1G and H). Moreover, the minor increase of Top2α expression by E6 is not due to protein stabilization caused by inhibition of proteasome degradation, illustrated in Fig. S1 that there was no significant difference in the expression of Top2α in MG132-treated E6 or E7 cells compared to the LXSN cell lines. The difference between E6 and the HPV genome on Top2α expression (see Fig. 1D) indicates that other HPV viral proteins and other regulatory mechanisms than transcription might participate in HPV-induced Top2α upregulation.

### Top2α upregulation is necessary for HPV replication

To assess whether Top2α acts on HPV replication, HPV31-positive CIN612 or HaCaT31 cells were transduced with either the control shRNA (TRC) or two Top2α-specific shRNA (shTop2α#1 or shTop2α#2) lentiviruses individually. The stable knockdown cell lines were selected by puromycin resistance. As shown in Fig. 2A, Top2α was efficiently knocked down in CIN612 cells. The Top2α-depleted cells were then differentiated in high calcium media and assayed with Southern blot analysis to examine the levels of HPV integrated genomes and episomes. Disruption of Top2α greatly suppressed HPV genome maintenance and amplification upon epithelial differentiation (Fig. 2B). A similar observation was also validated in HaCaT31 cells with stable Top2α knockdown (Fig. 2C). DNA lysates from the knockdown cells were extracted and assayed by DNA-qPCR, which showed that the viral DNA levels were significantly reduced in the silenced cells (Fig. 2D). The extrachromosomal fractions were further isolated for detection of HPV episomes by dot blot assay using digoxigenin-labeled HPV31 DNA probes (46). Fig. 2E illustrated that extrachromosomal HPV genomic DNA levels were markedly decreased by Top2α inhibition.

**Fig 2.**
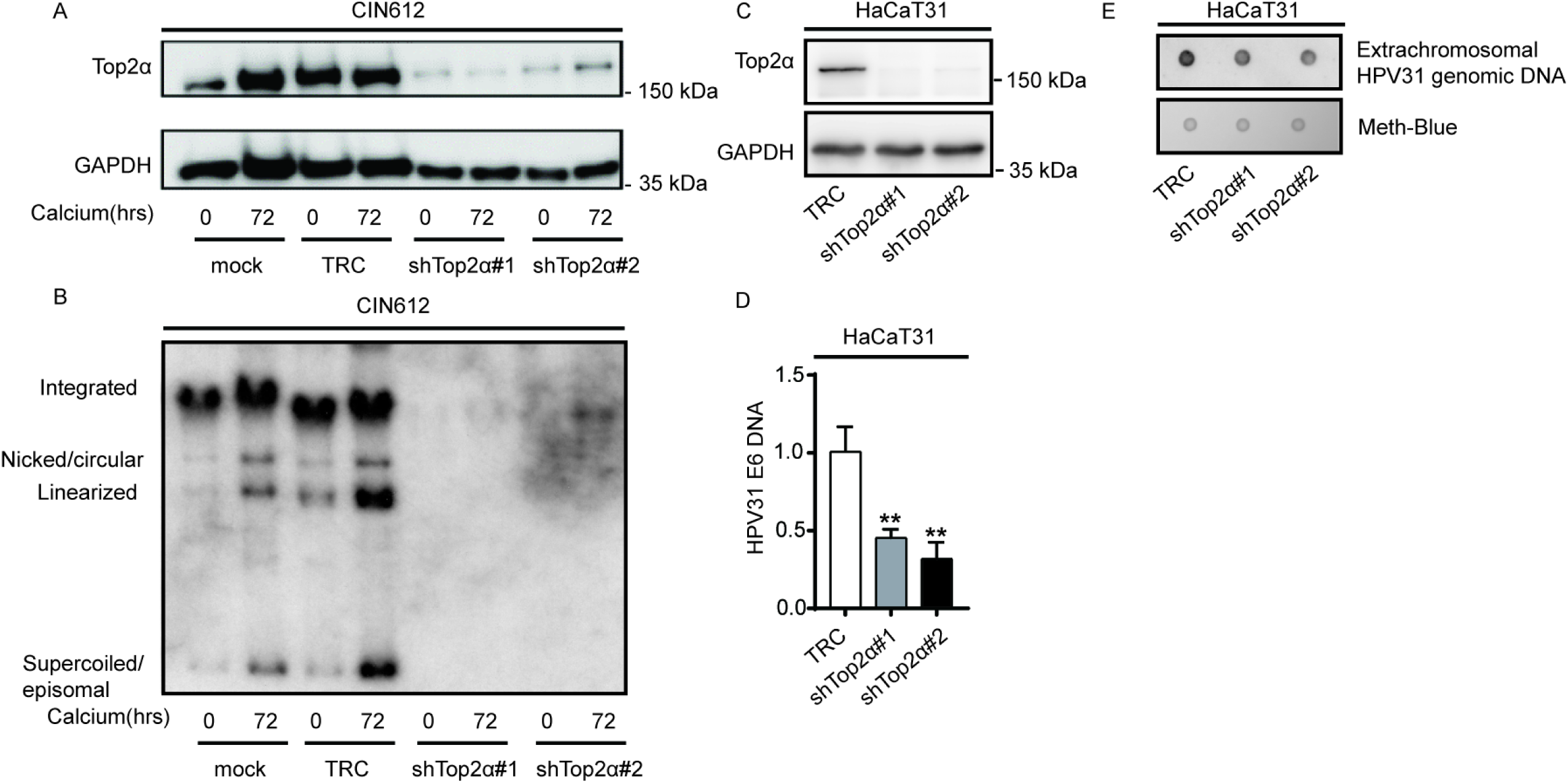
Top2α is necessary for HPV genome maintenance and amplification. (A) The Top2α knockdown efficiency was confirmed by western blot analysis in the CIN612 cell background. GAPDH served as the loading control. (B) DNA lysates in Top2α knockdown cells was harvested at 0 hours and 72 hours. Southern blot analysis was performed to examine viral episomes using the HPV31 genome as a probe. Integrated, nicked/circular, linearized, and supercoiled/episomal genomes are indicated. (C-E) HPV31-positive HaCaT31 cells were transduced with lentivirus containing control scramble shRNA sequences (TRC) or two shRNA sequences for Top2α (shTop2α#1 and shTop2α#2). The protein and DNA were extracted from HaCaT31-TRC, shTop2α#1, and shTop2α#2 cell lines. (C) Western blot analysis was carried out to compare Top2α, expression in HaCaT31-TRC, shTop2α#1, and shTop2α#2 cell lines, with GAPDH as the loading control. (D) DNA-qPCR was performed to measure the HPV31 E6 levels. Mitochondrial DNA (mtDNA) was used as the internal control for data normalization. The data were representative from three independent experiments. The statistical analysis was assayed by 2-tailed t test. Data are means ± standard errors. \*\**P*≤0.01. (E). Representative dot blot analysis of undifferentiated HPV31-containing keratinocytes with Top2α depletion.

### Loss of Top2α limits HPV replication in a DDR-independent manner

Given its canonical function in DDR, we tested whether Top2α promotes HPV replication in a DDR-dependent manner. The neutral comet assay was applied with HaCaT31 cells treated with the Top2α-specific inhibitor PluriSln#2 for detection of any changes of DNA double-stranded break (DSB). The DSB levels were quantified by measuring the percentage of DNA in the comet tail. The PluriSln#2 treatment did not increase the number of breaks, however, etoposide, a well-known inhibitor of both Top2α and Top2β, induced large amounts of DSB (Fig. 3A). This observation was confirmed using Top2α-silenced HaCaT31 cells. Increases in the broken fragments were minimally observed in the knockdown cells compared with the scramble (TRC) cells (Fig. 3B). The immunoblotting further showed that the expression of the DSB marker γH2AX was barely influenced by Top2α loss (Fig. 3C). Consistently, suppression of Top2α did not alter the expression or phosphorylation of DDR kinases such as ATM, ATR and DNA-PKcs, all of which are known to increase γH2AX levels (Fig. 3C). Altogether, these results show that Top2α does not impact the activation of DDR in these cells.

**Fig 3.**
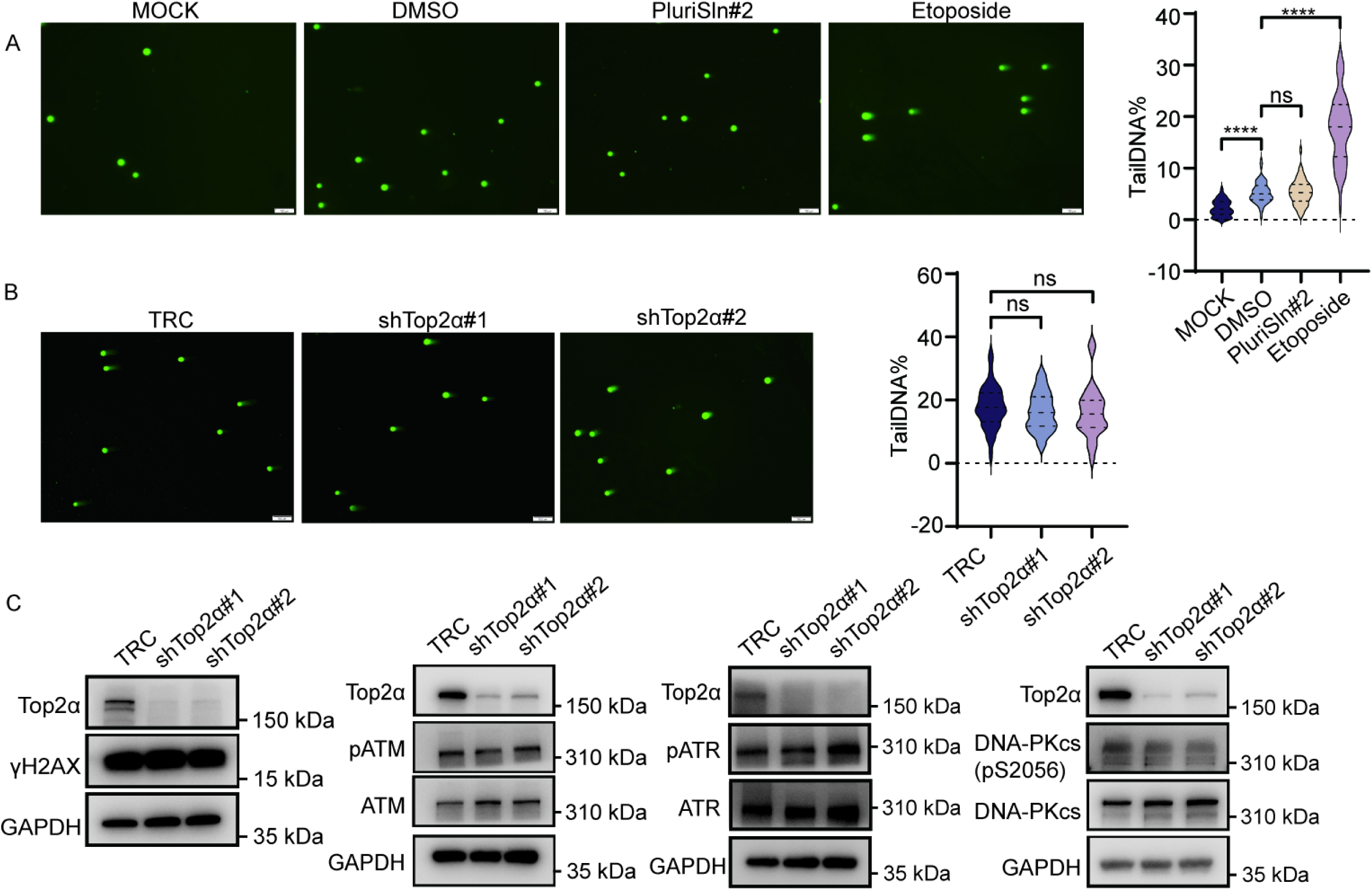
Top2α promotes HPV replication in DDR-independent pathway. (A) The comet assay in HaCaT31 cell lines that were treated with DMSO, PluriSln#2 (Top2α inhibitor, 10 μM), or etoposide (TOP2 inhibitor, 50 μM) for 6 hours. Quantitation of percent tail versus nucleoid body is shown in the violin plots (n=at least 50 number of nucleoids, ns=no significance, \*\*\*\**P*≤0.0001, Scale bar, 100 μm). (B) Comet assays for DNA break formation in HaCaT31 cells with TRC, shTop2α#1, and shTop2α#2. Quantitation of tail percentage versus nucleoid body is shown in the violin plots (n=at least 50 number of nucleoids, ns=no significance, Scale bar, 100 μm). (C) Western blot analysis was performed using antibodies to γH2AX, ATM, phosphorylated ATM, ATR, phosphorylated ATR, DNA-PKcs, and phosphorylated DNA-PKcs in Top2α-deficient cells. GAPDH served as the loading control. Data were representative of at least three independent biological repeats.

### Top2α regulates IL-6 and IL-8 secretion in HPV-positive cells

To explore downstream functions of Top2α in HPV-positive cells, an RNA-seq analysis was performed to compare the differential gene expression profiles between the Top2α-silenced HaCaT31 cells (shTop2α#1 or shTop2α#2) and the control cells. After RNA extraction and library preparation, the sequencing reads were processed. The data was normalized using the DESeq2 method. A cutoff standard of |log2FC|≥0.7, adj. p-value≤0.05, and p-value≤0.05 were applied. 192 differentially expressed genes (DEGs) were identified between shTop2α#1 and the control cells, and 559 ones were found between shTop2α#2 and the control cells. Of these, 117 DEGs were common in both comparisons (Fig. 4A, left panel), which were further analyzed using Metascape web-based portal for pathway enrichment analysis. As shown in Fig. 4A (right panel), these genes were enriched in several inflammation-related pathways including IL-4 and IL-13 signaling, inflammatory response, humoral immune response, and negative regulation of response to wounding. The expression of these inflammatory genes at the mRNA level was checked by RT-qPCR in the Top2α knockdown cells. Among the genes tested, there lack significant changes in the expression of IL-1B, IL-17, IL-18, TNF-alpha, CCL5, CCL20, CXCL9, CXCL10, CXCL11, or IFNB. However, IL-6 and IL-8 were increased whereas CCL2 was downregulated by Top2α inhibition (Fig. 4B).

**Fig 4.**
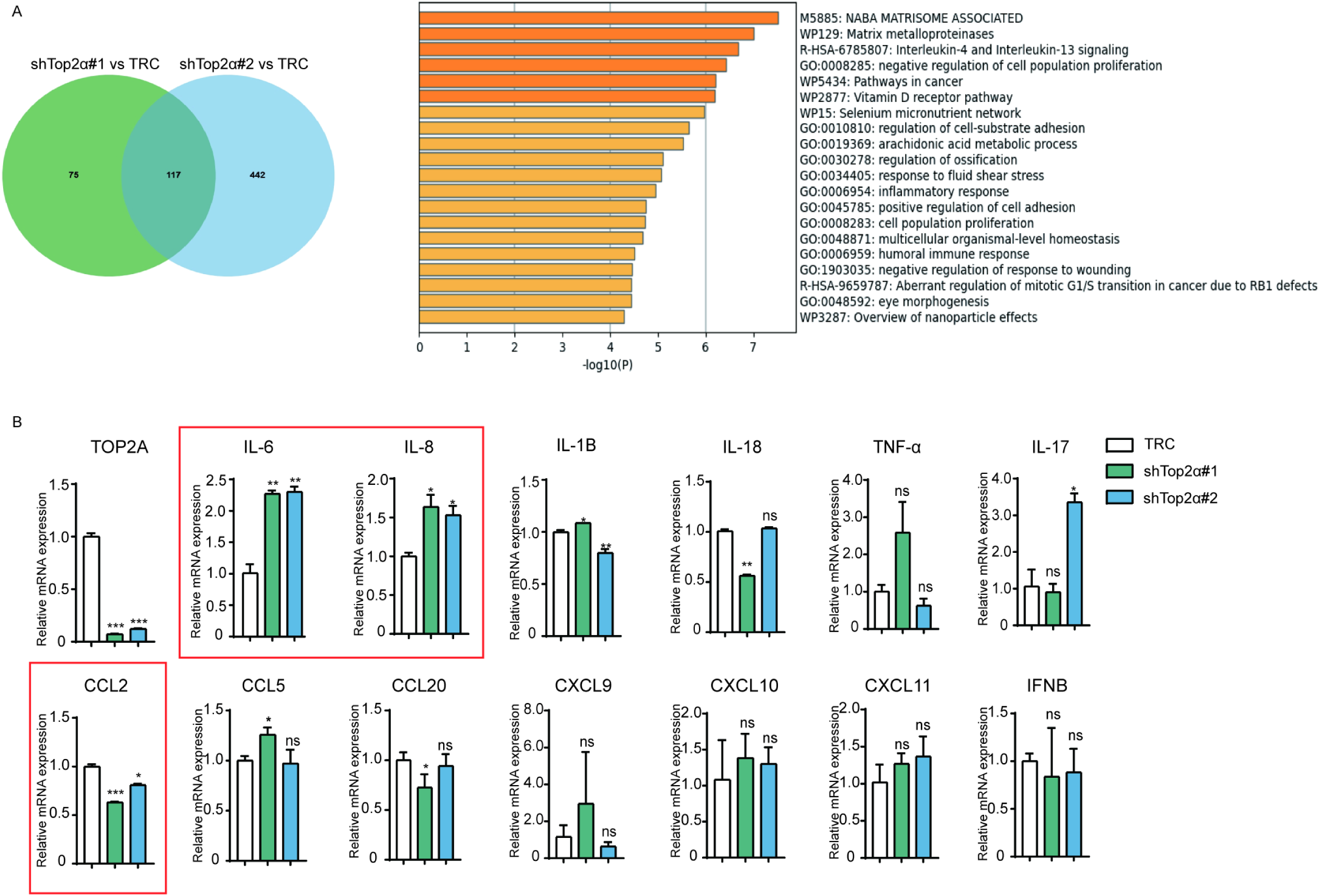
Top2α depletion might deregulate the expression of inflammatory genes in HPV-positive cells. (A) RNA-seq analysis was performed using HaCaT31 cells with Top2α knockdown. 117 differential expressed genes were identified as downstream targets of Top2α. Top20 pathways regulated by Top2α were listed with Metascape online analysis. (B) RT-qPCR analysis was conducted to assess the expression levels of TOP2A, IL-1B, IL-6, TNF-α, IL-8, IL-18, IL-17, CCL2, CCL5, CCL20, CXCL9, CXCL10, CXCL11, and IFNB in HaCaT31 cells with Top2α knockdown. GAPDH was used as the internal control for data normalization. Data are representative of at least three independent biological repeats. The statistical analysis was assayed by 2-tailed t test. \**P*≤0.05, \*\**P*≤0.01, \*\*\**P*≤0.001.

This complexity raised our interest in the role of IL-6, IL-8, and CCL2 during HPV life cycle. To assess their significance in cervical cancer development, the vaginal discharge was collected from normal, cervical intraepithelial neoplasia (CIN), and cervical cancer patients to measure the protein levels of these inflammatory factors using protein microarray technology (Table S1). The results showed that IL-6 levels were gradually increased in CIN and cervical cancer patients, compared to the normal group. IL-8 measurements were not interpretable as the levels were beyond the range of detection, while CCL2 production was not altered in cervical cancer (Fig. 5A). Given the previously reported role of IL-8 in HPV replication (35), the significance of IL-8 could not be ignored and we continued to include IL-8 as a potent target for Top2α studies. GEPIA datasets for cervical cancer patients were used to evaluate the prognostic values of IL-6, IL-8, and CCL2. High expression levels of IL-6 and IL-8 were associated with worse overall survival (OS) outcomes in cervical cancer patients (*P*≤0.05), but CCL2 was not related (*P*>0.05) (Fig. 5B). We continued to verify the total and the secreted protein levels of IL-6, IL-8, or CCL2 in the Top2α-depleted HaCaT31 cells. Different from the qPCR result (Fig. 4B), western blot showed that loss of Top2α had little, if any effect on the total protein levels of IL-6, IL-8, or CCL2 (Fig. 5C). Instead, the secreted levels of IL-6 and IL-8, but not CCL2, were significantly reduced in the knockdown cells, as determined by ELISA assay (Fig. 5D). These observations lead us to next investigate whether secreted IL-6 or IL-8 contributes to HPV replication.

**Fig 5.**
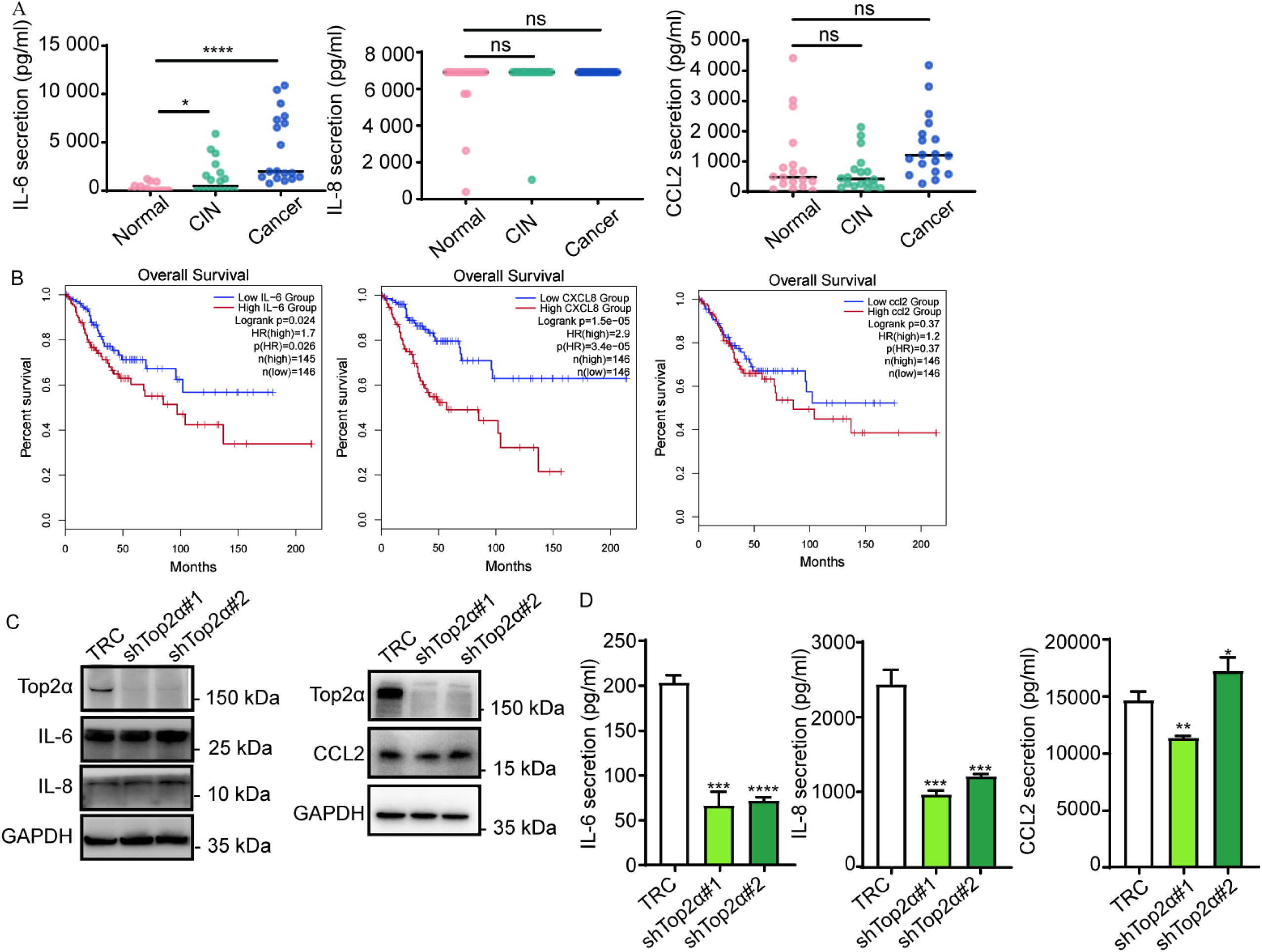
Top2α regulates IL-6 and IL-8 secretion in HPV-positive cells. (A) The protein microarray was utilized to measure the levels of IL-6, IL-8, and CCL2 in vaginal discharge samples from normal, cervical intraepithelial neoplasia (CIN), and cervical cancer patients. (B) Gene Expression Profiling Interactive Analysis (GEPIA) was performed to analyze the correlation of IL-6, IL-8, and CCL2 expression with the overall survival rate of cervical cancer patients. The red lines represent patients with high gene expression, and the blue lines with low gene expression. (C) Western blot demonstrating the expression of Top2α, IL-6, IL-8, and CCL2 proteins in the Top2α-depleted cell lines. GAPDH served as the loading control. (D) The levels of secreted IL-6, IL-8, and CCL2 outside Top2α knockdown cells were determined by ELISA assay. Data are representative of at least three independent biological repeats. The statistical analysis was assayed by 2-tailed t test. \**P*≤0.05, \*\**P*≤0.01, \*\*\**P*≤0.001, \*\*\*\**P*≤0.0001.

### IL-6/IL-8 promotes HPV replication

To test the effect of IL-6 on HPV genome maintenance, HaCaT31 cells were treated with the IL-6 neutralizing antibody (80 ng/mL) or the recombinant human IL-6 (10 ng/mL) for 72 hours. Extrachromosomal circular genomes were extracted from the treated cells using Hirt extraction method and analyzed by DNA-qPCR to detect HPV genes or genomes. Our data showed a significant decrease of HPV E6 expression by the treatment with the IL-6 neutralizing antibody, compared to the IgG control. Conversely, the addition of the IL-6 recombinant protein upregulated HPV E6 levels in the cells (Fig. 6A, left panel). We next examined HPV genome amplification in HaCaT31 cell lines cultured in high calcium media for 72 hours. High calcium treatment induced differentiation-dependent HPV genome amplification, represented by a significant increase of E6 gene levels. This amplification was greatly reduced by the treatment of the IL-6 neutralizing antibody (Fig. 6A, right panel). In contrast, IL-6 treatment resulted in increased E6 levels compared to IgG control or empty treatment. These data demonstrated that IL-6 is important for HPV maintenance and amplification (Fig. 6B). The differentiation marker involucrin expression was visualized by western blot in differentiating HPV31-positive cells, which were induced by high calcium culture and not affected by the treatment of IL-6 or the neutralizing antibody (Fig. 6C). The similar approaches found that IL-8 neutralizing antibody suppressed viral gene expression and significantly inhibited the viral replication in undifferentiated cells or differentiated cells cultured in high-calcium media for 72 hours. Contrasted with the IL-8 neutralizing antibody, IL-8 treatment promoted HPV31 gene expression and genome amplification (Fig. 6D). In contrast, CCL2 treatment failed to augment HPV E6 gene expression or amplification (Fig. 6E).

**Fig 6.**
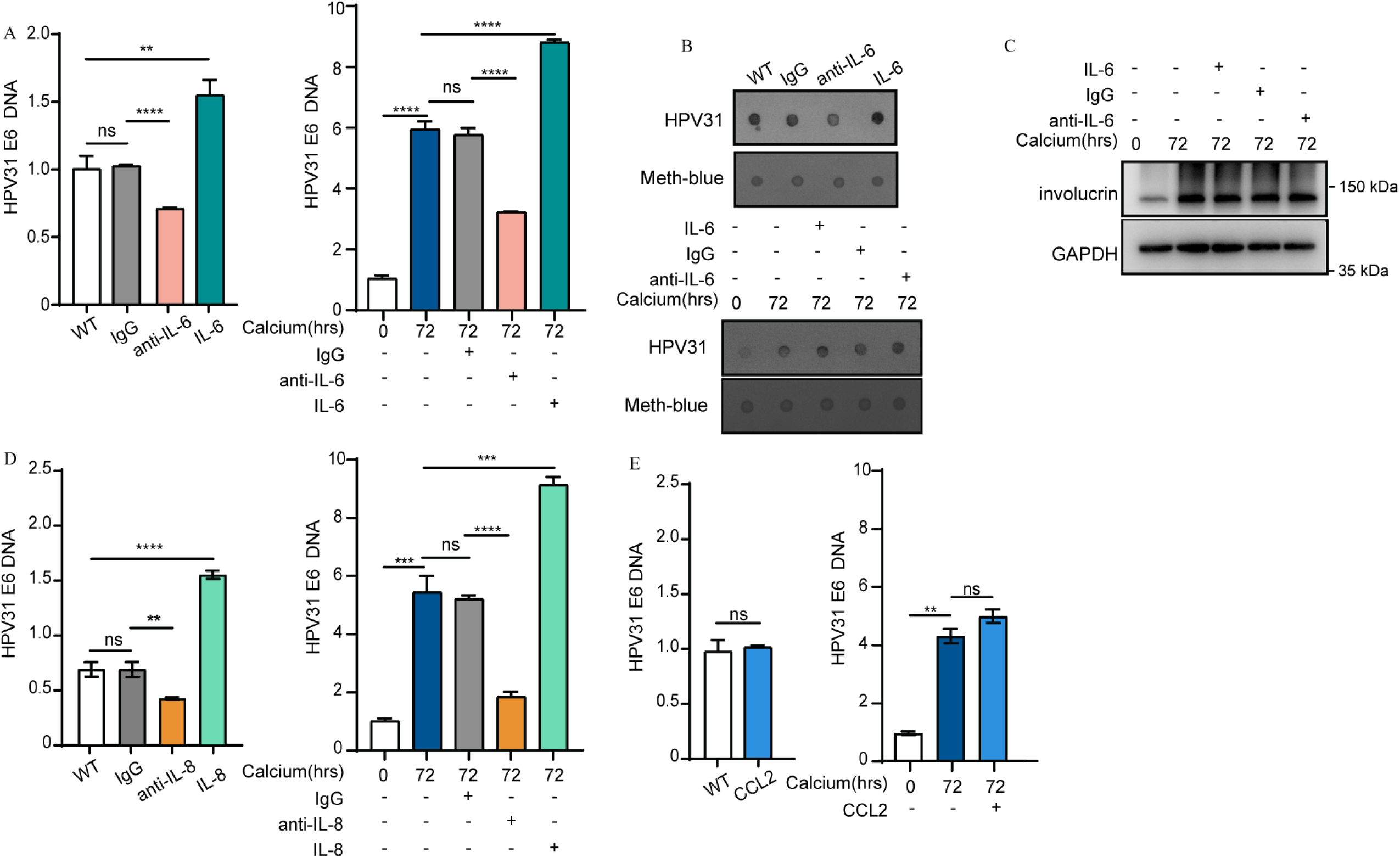
IL-6/IL-8 promotes HPV replication. (A) DNA-qPCR was performed to detect HPV episomes in the undifferentiated or differentiated HaCaT31 cells that treated with IL-6 (10 ng/mL) or an anti-IL-6 neutralizing antibody (80 ng/mL) for 72 hours. Shown is the relative HPV DNA levels in the treated cells normalized to the untreated cells. (B) Dot blot analysis was applied with the HPV31-specific probe to measure HPV episomes. Meth-blue served as the loading control. (C) Western blot demonstrating the expression of involucrin proteins with GAPDH as the loading control. (D) The DNA was extracted from undifferentiated or differentiated HaCaT31 cells that treated with IL-8 (10 ng/mL) and an anti-IL-8 neutralizing antibody (50 μg/mL) for 72 hours. (E) The DNA was extracted from undifferentiated or differentiated HaCaT31 cells that treated with CCL2 (100 ng/mL) for 72 hours. Data are exhibited as means ± SEM, and are representative of at least three biological independent repeats. The statistical analysis was assayed by 2-tailed t test. ns=no significance, \*\**P*≤0.01, \*\*\**P*≤0.001, \*\*\*\**P*≤ 0.0001.

**TLR4 participates in secretion of IL-6 and IL-8 regulated by Top2α**

We next searched our RNA-seq data for the inflammation-related regulators of cytokine/chemokine expression or secretion to investigate the details of Top2α-dependent IL-6/IL-8 secretion. A heatmap of the gene profiles related to inflammatory pathways was demonstrated in Fig. 7A. Among them, FN1, TLR4, and PLA2G2A were significantly decreased in Top2α silencing cells. Previous studies have indicated that TLR4 regulates the secretion of IL-6 and IL-8 (47). FN1 encodes fibronectins that bind to collagen or fibrin to function in cell adhesion and cell motility, while PLA2G2A is a member of secretory phospholipase A2 with undefined functions. Based on these findings, we selected TLR4 as the proposed target for further investigation. TLR4 expression at mRNA or protein levels were then examined in Top2α-depleted cells, and our data showed that Top2α silencing significantly reduced TLR4 expression (Fig. 7B and C). To further check whether Top2α regulates TLR4 transcription, the luciferase assay and ChIP-qPCR assay were applied with Top2α-silencing cells. The results showed that loss of Top2α significantly decreased the activities of the TLR4 promoter (Fig. 7D). To explore whether Top2α affects TLR4 transcription by directly binding to the TLR4 gene promoter, three sets of ChIP primers were selected by Primer3Plus online tool to target three regions to which Top2α likely binds (Fig. 7E). Among these three regions, the -387 — -247bp region contains the binding sequence of PU.1, which has been indicated that regulates the TLR4 gene expression (48). ChIP-qPCR assay demonstrated that Top2α ubiquitously bound to TLR4 promoter region at these binding regions (Fig. 7F). To explore the effects of TLR4 on IL-6/IL-8 secretion and HPV replication, we treated HaCaT31 cells with TAK-242, a selective inhibitor of the TLR4 pathway. Fig. 7G show significant reduction of secreted IL-6 or IL-8 levels in the cell culture, with a clear dose-dependent effect of TAK-242. Moreover, TAK-242 treatment resulted in a suppression of HPV genome and E6 expression (Fig. 7H and I). Taken together, Top2α transcriptionally regulates TLR4 expression to promote IL-6/IL-8 secretion and HPV replication.

**Fig 7.**
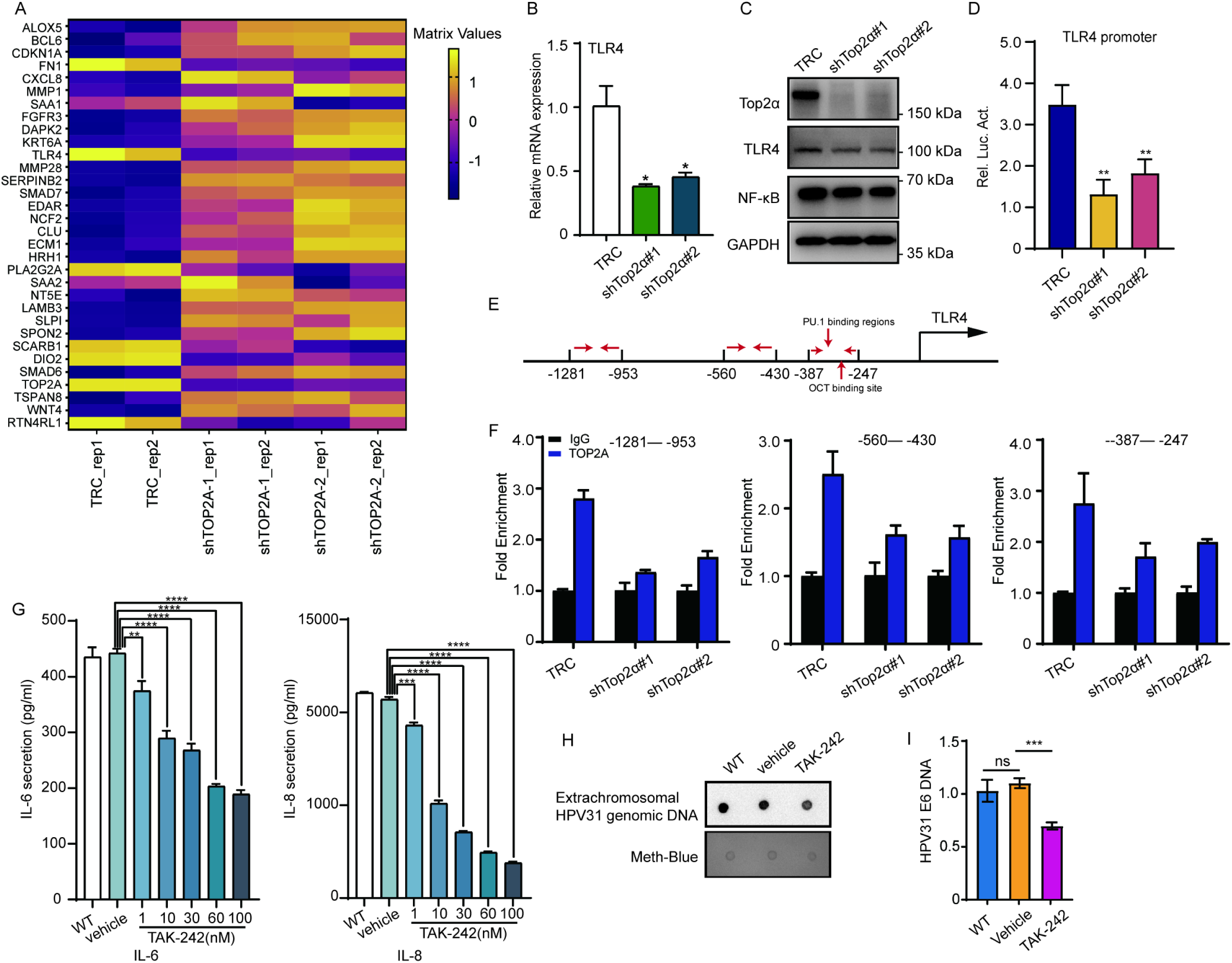
TLR4 is a key mediator for Top2α-dependent IL-6/IL-8 secretion and HPV replication. (A) Heat maps of inflammatory regulatory genes regulated by Top2α knockdown. TRC_rep1 and TRC_rep_2 are replicates of scramble HaCaT31 cells, while shTop2α#1 (rep1 and rep2) and shTop2α#2 (rep1 and rep2) are replicates of HaCaT31 cells with Top2α knockdown respectively. (B) RT-qPCR was performed to detect TLR4 mRNA levels in Top2α knockdown cells. (C) Western blot analysis was performed using antibodies of Top2α, TLR4, and NF-κB, with GAPDH as the loading control. (D) The TLR4 promoter plasmid were transfected into HaCaT31-TRC, shTop2α#1, and shTop2α#2 cell lines. A pRL-TK plasmid was transfected to normalize the transfection efficiency data. Luciferase activity was measured in cell lysates for 48 hours post-transfection. Data are exhibited as means ± SEM, and are representative of at least two biological independent repeats. \*\**P*≤0.01. (E and F) ChIP-qPCR was performed in Top2α-depleted cells to identify the binding of Top2α on the TLR4 promoter regions. IgG served as an antibody control. (G) IL-6 and IL-8 were quantified in the supernatants from outside of HaCaT31 cells with TAK-242 treatment of indicated concentrations. \*\**P*≤0.01, \*\*\**P*≤0.001, \*\*\*\**P*≤0.0001. (H and I) The viral DNAs extracted from HaCaT31 cells treated with DMSO or TAK-242 (10 nM) for 72 hours were visualized by dot blot or DNA-qPCR. Data are representative of three biological independent repeats. ns=no significance, \*\*\**P*≤0.001.

## DISCUSSION

Our work demonstrates that Top2α functions as a pivotal regulator through a novel DDR-independent mechanism to facilitate the viral genome maintenance and amplification. This action is achieved by regulating the secretion of key inflammatory cytokines IL-6 and IL-8. TLR4 serves as a key mediator in this regulation. Inhibition of either Top2α or TLR4 effectively reduces HPV replication, demonstrating their central involvement in HPV life cycle. Moreover, treatment of HPV-positive cells with IL6/IL-8 neutralizing antibodies further confirms that these cytokines are essential for HPV genome amplification. Our findings reveal a complex relationship between Top2α, TLR4 and cytokine secretion, suggesting that HPV manipulates host inflammatory pathways to create a tumor-promoting microenvironment conducive to the viral replication (Fig. 8). These insights offer new perspectives on HPV pathogenesis and open avenues for novel therapeutic interventions aimed at disrupting the Top2α-TLR4-cytokine axis in HPV-associated cancers.

**Fig 8.**
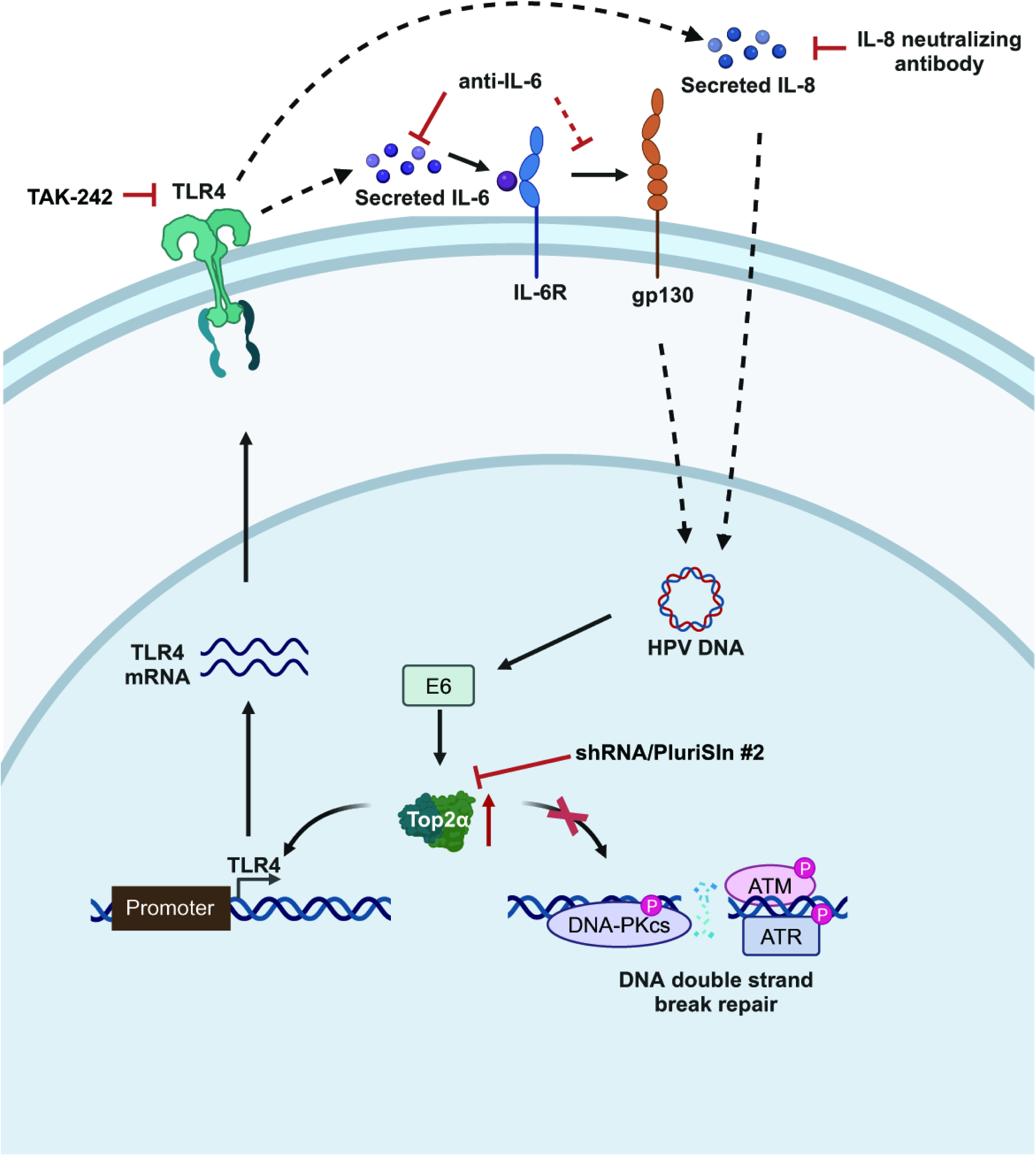
A schematic illustration for manipulation of Top2α-TLR4 signaling by HPV to regulate IL-6/IL-8 secretion and HPV replication. HPV induces Top2α, partially through E6, which then regulates TLR4 transcription by directly binding to TLR4 promoter region. The elevated TLR4 signaling facilitates the secretion of IL-6/IL-8 that in turn promotes HPV replication. Inhibition of Top2α by shRNA approaches, suppression of TLR4 by TAK-242, and neutralization of IL-6/IL-8 significantly diminishes HPV replication. The image is drawn using Biorender software.

DDR proteins TopBP1 and Top2β (19, 20) promote HPV genome amplification and maintenance. However, Top2α functions differently from its homolog Top2β to promote HPV replication, as our findings show that it is independent of DDR activation or DSB formation. The structures of TOP2A and TOP2B when complexed with DNA and etoposide were aligned and the analysis showed no significant differences (Fig S2). However, the C-terminal domains (CTDs) of Top2α and Top2β differ structurally, which likely accounts for their functional divergence. Despite the overall similarity in structure, the CTDs of Top2α and Top2β share only 34% amino acid identity (Fig S2). Notably, Top2α’s CTD lacks nine phosphorylation sites and one SUMOylation site found in Top2β’s CTD, which are implicated to be critical for Top2β’s roles in DNA-binding, decatenation, or interaction with other proteins (49, 50). The unique structure of Top2α CTD positions Top2α as a non-redundant regulator of HPV replication, which acts through modulation of host inflammation rather than via canonical DDR mechanisms. Consistent with Top2α, other two other DDR proteins, TopBP1 and ATR, also contribute to the regulation inflammatory gene expression in HPV-positive cells, which fits into the idea that HPV walks on the line between DDR and inflammation.

Previous studies have shown that DDR mingles with inflammation as a host defense against exogenous stimuli or endogenous stresses (51, 52). Our work demonstrates that Top2α mediates TLR4 to effectively facilitate secretion of IL-6 and IL-8, which in turn promote HPV replication. Both IL-6 and IL-8 play critical roles in RNA virus replication (53, 54) or establishment of tumor microenvironment (55, 56). IL-6 has been reported to exert either pro- or anti-viral effects depending on the context (57, 58), and our data confirm its role in promoting HPV genome amplification upon epithelial differentiation (Fig. 6A). This finding is consistent with prior evidence of linking IL-6/STAT3 signaling to HPV-associated cervical cancer progression (36). Moreover, secreted IL-6 enhances radiosensitivity of HPV-related cancers via inducing M1 macrophage polarization (59), indicating its potential as either a biomarker or a target of adjunctive therapies for HPV-related malignancies. Similarly, we have shown that IL-8 is critical to HPV gene expression (Fig. 6C), consistent with the Clayberger group’s finding that neutralization of IL-8 suppresses HPV replication upon epithelial differentiation (35). In addition to HPV, antibody neutralization of endogenous IL-8 inhibits HCV replication (60), further supporting the broader importance of IL-8 in viral replication.

HPVs often use their oncoproteins to regulate DDR signaling. Here we tested whether E6 or E7 transcriptionally regulates Top2α expression or whether they effect its stability through proteasome-dependent mechanism. We observed that HPV oncoprotein E6, but not E7, modestly upregulates Top2α mRNA expression levels (Fig. 1E and 1F), indicating that the regulation occurs at the post-transcriptional level, likely through protein stabilization or microRNA modulation. We found no evidence that the proteasomal degradation is involved as Top2α expression was unaffected in E6-/E7-overexpressing cells treated with MG132 (Fig. S1). This rules out the possibility of proteasome-dependent degradation. Recent studies indicate that E6 downregulates miR-320a to increase Top2α expression (61). Additionally, predictions from miRDB and TargetScan databases indicate that other microRNAs such as miR-139-5p (62), miR-345-5p (63) or miR-144-3p (chosen by the highest score), may contribute to the regulation of Top2α expression, highlighting a potential area for further exploration. Moreover, western blot analysis showed E6 modestly increases Top2α protein expression, which suggests that other viral proteins might contribute to induction of Top2α. Overall, these possibilities showed that the regulation of Top2α expression by HPV is more complicated than initially thought, warranting further investigation into the underlying mechanisms.

To our knowledge, this is the first evidence, at least in HPV field, that the DDR protein can regulate cytokine secretion to shape the host microenvironment, thereby facilitating HPV replication. TLR4 has dual roles in both inflammation and HPV replication, with our findings highlighting its importance in the viral life cycle. Our study underscores the potential of targeting Top2α, TLR4, and the associated inflammatory cytokines IL-6 and IL-8 as part of a multipronged approach to treating HPV-related diseases. IL-6 inhibitors or neutralizing antibodies, which are currently being explored in oncology, could be repurposed to disrupt the tumor-promoting microenvironment in HPV-positive cancers. Similarly, blocking TLR4 signaling with inhibitors like TAK-242 offers a novel therapeutic avenue to reduce both viral replication and inflammation-driven carcinogenesis. Collectively, our study unveils a previously unrecognized axis critical to HPV replication and development of HPV-related cancers.

## MATERIALS AND METHODS

### Clinical tissue samples

The human cervical biopsy tissue samples analyzed in this study was approved by the Institutional Review Board (IRB) of Women’s Hospital of Zhejiang University (IRB-20220235-R). The samples were collected from the Women’s Hospital, School of Medicine, Zhejiang University (Zhejiang Province, China). Prior to sample collection, each subject who underwent surgical resections signed a written informed permission form.

### RNA extraction and Quantitative Real-time PCR (qPCR)

Total RNA was isolated from the cultured cells using TRIzol (Invitrogen, #15596018CN) according to the manufacturer’s instructions. 1 μg of total RNA was then reversely transcribed into cDNA products using ABScript Neo RT Master Mix for qPCR with gDNA remover (ABclonal, #RK20433). qPCR was performed using 2× Universal SYBR Green Fast qPCR Mix (ABclonal, #RK21203) on the LightCycler 480 real-time PCR instrument. The 2^-ΔΔCT^ method was used to calculate the relative levels of target genes, normalized to either GAPDH or EEF1A1 levels. The representative data are from three independent experiments. Significance was determined using Student’s t-test and a *P* value of < 0.05 was considered significant. The primers used for RT‒qPCR are listed in Table S2.

### Cell lines and cell culture

Human foreskin keratinocytes (HFKs) were obtained from the American Type Culture Collection. Human keratinocytes maintaining HPV16 and HPV31 episomes were generated by transfection of HFK with viral genomes as previously described (64). The spontaneously immortalized human keratinocyte cell line HaCaT, along with NIH 3T3 J2 fibroblasts and PT67 cells, were sourced from Procell Life Science and Technology Co., Ltd (Wuhan, China). HPV genome-expressing cells or E6- or E7-expressing HaCaT cell lines, were selected using G418 after transfection with the HPV genome or infection with retroviruses, as previously described (65).

All HFK and HPV-positive cells were maintained in E-medium supplemented with 5 ng/mL mouse epidermal growth factor (Sigma, #SRP3196). To induce differentiation, the cells were co-cultured with mitomycin C-treated NIH 3T3 J2 fibroblasts in keratinocyte basal medium supplemented with growth supplements. The cells were then switched to keratinocyte basal medium (without supplements) containing 1.5 mM CaCl_2_ for up to 72 hours.

HaCaT, HaCaT16, HaCaT31, E6- and E7-expressing cells were cultured in Minimum Essential Medium α (MEM α, Gibco), supplemented with 100 U/mL penicillin, 100 μg/mL streptomycin, and 10% heat-inactivated fetal bovine serum. Cells were incubated at 37°C in a 5% CO_2_ atmosphere. To induce differentiation, cells were cultured in Medium 154 (Gibco, #M154500) containing 0.03 mM CaCl_2_ with growth supplements for at least 24 hours, after which they were switched to Medium 154 containing 1.5 mM CaCl_2_ without growth supplements for up to 72 hours.

### Lentiviral particle production and transduction

Lentiviruses were produced as previously described(7). Two Top2α-specific shRNA constructs or a non-targeting shRNA control construct were transfected into 293T cells, along with the packaging plasmid psPAX2 and the envelope plasmid pMD2.G, to generate lentiviral particles individually. Each of these plasmids (2 μg) was co-transfected with 0.9 μg of psPAX2, 0.9 μg of shRNA, and 0.2 μg of pMD2.G plasmids into 293T cells using polyethyleneimine (PEI). Supernatants containing lentiviruses were harvested 48 hours post-transfection, filtered sterilized, and stored at −80°C until use. CIN612 and HaCaT31 cells were transduced with 0.5 mL of viral supernatants. The supernatant contained either the scramble or Top2α shRNA lentivirus particles, along with 10 μg/mL polybrene, and was incubated at 37°C for 12 hours. The cells were maintained in fresh media for an additional 48 hours. Knockdown of Top2α was confirmed through western blot analysis.

### Reagents and antibodies

IL-6 (MCE, #HY-P7044), IL-8 (MCE, #HY-P7379), CCL2 (MCE, #HY-P7237), IL-6 neutralizing antibody (SinoBiological, #10395-MHK23), anti-IL-8 antibody (Selleck, #A2524). Anti-TOP2A (Rabbit, Proteintech, #20233-1-AP, 1:200), anti-TOP2A (Mouse, Proteintech, #66541-1-Ig, 1:2500), anti-GAPDH (Mouse, Proteintech, #60004-1-Ig, 1:5000), anti-γH2AX (Mouse, Abmart, #M63324M, 1:2000), anti-NF-κB p65 (Mouse, Abmart, #T55034F, 1:1000), anti-phospho-ATM (Rabbit, Cell Signaling, #13050S, 1:1000), anti-phospho-ATR (Rabbit, Cell Signaling, #2853S, 1:1000), anti-ATM antibody (Rabbit, Cell Signaling, # 2873S, 1:1000), anti-ATR (Rabbit, Cell Signaling, #2790S, 1:1000), anti-DNA-PKcs (Mouse, Zenbio, #200618-6D1, 1:1000), anti-phospho-DNA-PKcs (Rabbit, Abmart, # PA5877S, 1:1000), anti-cytokeratin 10 (Rabbit, Zenbio, #R381139, 1:1000), anti-TLR4 (Rabbit, Abclonal, #A5258, 1:1000), anti-IL-6 (Rabbit, Abclonal, #A0286, 1:1000), anti-IL-8 (Rabbit, Wanleibio, #WL03074, 1:1000), anti-MCP-1 (Rabbit, Wanleibio, #WL02966, 1:1000), anti-p53 (Rabbit, Abclonal, #A26037, 1:1000).

### Western blot assay

The cells were lysed in RIPA lysis buffer (Solarbio, #R0010) supplemented with protease and phosphatase inhibitors for 30 min on ice and centrifuged at 12,000 rpm for 10 min at 4°C. After SDS-PAGE, the proteins were transferred to the PVDF membrane. The membrane was saturated in 5% skim milk for 1 hour and incubated overnight at 4°C with primary antibodies. The membrane was washed three times with tris buffered saline tween 20 (TBST) and was then incubated with secondary antibodies for 1 hour at room temperature. After three additional washes, protein bands were visualized by an ECL detection kit (Life-iLab, #AP34L024) and quantified using Chemiscope S6 (Shanghai, China).

### Immunohistochemistry (IHC)

The 4.0-µm thick sections of tumor tissues were deparaffinized and rehydrated. Heat-induced antigen retrieval was performed using 1× EDTA antigen retrieval solution (pH 9.0) (Solarbio, #C1034). Endogenous peroxidase activity was blocked using enhanced endogenous peroxidase blocking buffer (Beyotime, #P0100B). The slides were then washed and incubated in blocking buffer (Beyotime, #P0260) at 37°C before being incubated overnight at 4°C with anti-TOP2A (Rabbit, Proteintech, #20233-1-AP, 1:50). For signal amplification, a Horseradish peroxidase (HRP)-linked secondary antibody (ZSGB-BIO, #PV-6000) was used. 3, 3’ N-Diaminobenzidine Tertrahydrochloride (DAB) (ZSGB-BIO, #PV-8000) was applied for color development to detect Top2α. The reaction was stopped by washing with distilled water. The slides were then counterstained with hematoxylin, dehydrated, cleared, and sealed with neutral balata. Observations were made under an OLYMPUS BX53 microscope.

### TLR4 promoter reporters and dual-luciferase reporter assay

Human TLR4 promoter reporter plasmid of 2000bp was cloned into the pGL3-Basic vector upstream of the luciferase reporter gene. TLR4 promoter reporter plasmid was transfected into Top2α-silencing cells using Attractene Transfection Reagent (Qiagen, #301005) for 48 hours. After transfection, cells were lysed with 100 μL of lysis buffer and 20 μL of cell lysate was subjected to a dual luciferase reporter assay (Promega, #E1910). The firefly luciferase signals and Renilla luciferase signals were detected by an EnVision Mutilabel Reader. The luciferase reporter activity was calculated and all values were expressed as fold induction relative to the basal activity.

### Chromatin immunoprecipitation (ChIP)-qPCR

Approximately HaCaT31 (1 × 10^7^) cells were fixed in 1% formaldehyde (Cell Signaling Technology, #12606) for 10 min at room temperature, after which 125 mM glycine was added and left for 5 min to terminate the crosslinking reaction. The cells were then treated with ChIP lysis buffer 1 (50 mM HEPES-KOH, pH 7.5, 140 mM NaCl, 1 mM EDTA, 10% glycerol, 0.5% NP-40, 0.25% Triton X-100, and 1× protease inhibitors). The nuclear pellet was resuspended and rocked on a platform rocker at 4°C for 10 min, followed by centrifugation at 1000×g for 5 min at 4°C. Next, the pellet was treated with ChIP lysis buffer 2 (10 mM Tris-HCl, pH 8.0, 200 mM NaCl, 1 mM EDTA, 0.5 mM EGTA, and 1x protease inhibitors), resuspended again, rocked at 4°C for 10 min, and collected by centrifugation at 1000×g for 5 min at 4°C. The pellet was resuspended in lysis buffer 3 (10 mM Tris-HCl, pH 8.0, 1 mM EDTA, 0.5 mM EGTA, 0.1% SDS, and 1x protease inhibitors). The chromatin was sonicated to achieve an average DNA fragment length of 200–1000 bp by 30 seconds ON, 30 seconds OFF for 15 cycles using bioruptor plus (Diagenode) and then incubated 4 μg of rabbit anti-TOP2A (Proteintech, #20233-1-AP) or normal rabbit IgG (Proteintech, #30000-0-AP) antibodies at 4°C overnight for chromatin immunoprecipitation. DNA were extracted by the phenol‒chloroform method. Then, qPCR was utilized to quantify the immunoprecipitated DNA, and the data were normalized to the IgG antibody. Data are expressed as fold of IgG control ± SEM. The primers used for ChIP‒qPCR are listed in Table S3.

### Viral DNA extraction and dot blot hybridization

HPV DNA was extracted from cells as previously described (46). Briefly, two million cells were collected, resuspended, lysed, and precipitated. The cells were resuspended in 200 μL Solution 1 and transferred to a microfuge tube. Then, 400 μL Solution 2 was added to lyse the cells until the solution became clear and viscous. The mixture was then incubated for 5 min on ice. Following this, 300 μL of Solution 3 was added, and the solution was mixed until a white precipitate formed. This mixture was incubated on ice for additional 10 min. Subsequently, the sample was centrifuged at ≥ 16,000× g for 5 min at 4°C to pellet DNA and proteins. The supernatant was carefully transferred to a new tube, and isopropanol was added to precipitate nucleic acids. After centrifuging again to precipitate low molecular weight nucleic acids, the pellet was resuspended in Hirt buffer and incubated at 37°C for 30 min, followed by incubation at 50°C for another 30 min. Prepare a column in phase lock gel-heavy tubes and add the sample. Phenol:chloroform:isoamylalcohol was added, mixed, and centrifuged. This extraction process was repeated. Subsequently, chloroform:isoamylalcohol was added to the upper phase, mixed, and centrifuged again. The upper phase was transferred to a new tube, and DNA was precipitated with sodium acetate and ethanol, followed by incubation on ice for 30 min. The sample was then centrifuged to pellet the DNA, after which the supernatant was removed, and the pellet was washed with 70% ethanol and dried. Finally, the pellet was resuspended in TE buffer containing RNase A and incubated at 68°C for 20 min.

Equal amounts of DNA were loaded onto a positively charged nylon membrane and 1 mol/L NaOH was used for DNA degeneration. The membrane was prehybridized for 1 hour, after which the DIG-labeled probe was added to the buffer used for the prehybridization stage to hybridize at 42°C overnight in a hybridization incubator. The membrane was then washed twice with the primary wash buffer for 5 min at room temperature, followed by two washes with the secondary wash buffer for 15 min at 68°C. It was blocked with blocking solution (Roche, # 11096176001) for 2 hours at room temperature, and then incubated with the antibody solution (anti-Digoxigenin-alkaline phosphatase antibody) for 30 min at room temperature. The excess secondary wash buffer was drained from the blots and the membrane was transferred to the detection buffer (0.1 M Tris-HCl, 0.1 M NaCl, pH 9.5) for 5 min at room temperature. The detection reagent was applied to the blots for 5 min and quantified using the instrument (Chemiscope S6).

### HPV DNA content PCR

HPV DNA-qPCR was performed as previously described (66). The extraction of viral DNA extraction and qPCR were performed using 2X Universal SYBR Green Fast qPCR Mix on the LightCycler 480 real-time PCR instrument. The HPV31 DNA levels were measured using primers to HPV31 E6. HPV DNA was normalized to mtDNA using the primers. The primers used for DNA‒qPCR are listed in Table S4.

### Southern blot

DNA isolation and Southern blotting were performed as previously described (67). Briefly, cells were collected in Southern buffer (400 mM NaCl, 10 mM Tris, pH 7.5, and 10 mM EDTA). The cell membranes were disrupted by adding 30 μL of 20% SDS, followed by incubation overnight at 37°C with 15 μL of a 10 mg/mL proteinase K solution to digest proteins. The DNA was extracted using a phenol-chloroform procedure and then precipitated with a mixture of sodium acetate and ethanol. The purified DNA samples were subjected to restriction enzyme digestion with BamHI, which does not cut the HPV31 genome. The DNA fragments were resolved by electrophoresis on a 0.8% agarose gel and then transferred onto a positively charged nylon membrane. Hybridization was performed using ^32^P-labeled HPV31 genome as a probe.

### Enzyme-Linked Immunosorbent Assay (ELISA)

IL-6 and IL-8 levels in the supernatants were quantified using commercially available ELISA kits, following the manufacturer’s protocol (Abclonal, #RK00004, #RK00011). The diluted samples and standards were added to the wells and incubated at 37°C for 2 hours. After incubation, the plates were washed, and a biotin-conjugated antibody was added, followed by incubation at 37°C for 60 min. The plates were washed again, and streptavidin-HRP antibody was added to each well and incubated at 37°C for 30 min. Following another wash, the tetramethyl-benzidine (TMB) substrate was added in each well. After a 15-20 minutes incubation at 37°C, a stop solution was added, and the optical densities were read at 450 nm and 570 nm wavelengths. All experiments were performed in triplicate. The results for IL-6 and IL-8 were expressed as concentrations in pg/mL.

The concentration of CCL2 in the supernatants was determined by the Human CCL2 ELISA Kit (UpingBio, #SYP-H0576). The diluted samples, standards and biotin-conjugated antibody were added to the wells and incubated at 37°C for 60 min. Then, the wells were washed and HRP-conjugate reagent was added and incubated at 37°C for 20 min. The wells were washed and color developing reagent was added in each well. After 15 min incubation at 37°C, stop solution was added and the optical densities were read at a wavelength of 450 nm. All experiments were performed in triplicates. The results of CCL2 were expressed as concentrations in pg/mL.

### Neutral Comet assay

According to CometAssay^®^ ES II electrophoresis system (Trevigen), cells were trypsinized, counted, and embedded in LMAgarose on two-well slides at a density of 1000 cells per well, then lysed overnight at 4°C. The cells were immersed in 1× neutral electrophoresis buffer for 30 min at 4°C, after which electrophoresis was performed at 21 V for 45 min. Following electrophoresis, the cells were immersed in DNA precipitation solution for 30 min at room temperature, followed by immersion in 70% ethanol for another 30 min at room temperature. The samples were dried at 37°C for 15 min and each sample was stained using 100 μL of diluted SYBR^®^ Gold (Invitrogen, #S11494) for 30 min in the dark at room temperature. Cells were allowed to dry completely at 37°C and were visualized using an OLYMPUS BX53 microscope. The TailDNA% was quantified using CASP software analysis.

### RNA sequencing

The Illumina sequencing was performed at Novogene Bioinformatics Technology Co., Ltd., in Beijing, China. The cells were washed with PBS, flash frozen on dry ice, and the RNA was harvested using Trizol reagent. The construction of sequencing libraries was performed using the Illumina TruSeq RNA Sample Prep Kit (#FC-122-1001), using 1 μg of total RNA in accordance with the standard protocols established by Novogene. Raw counts were used to assess the quality of raw and post-alignment sequencing reads. Subsequently, the reads were aligned to the human GRCh38 genome model using Hisat2 v2.0.5. The identification of differentially expressed genes (DEGs) was conducted using the DESeq2 method, comparing the knockdown groups with the control groups. Each group included duplicate biological specimens. A pathway enrichment analysis was performed using Metascape. The identified significant DEGs were those that met the criteria of adjusted *P* (FDR) <0.05 and |log2fold change (FC)|≥0.7. The original data of the RNA-seq was uploaded to the GEO DataSets (https://www.ncbi.nlm.nih.gov/geo/query/acc.cgi?acc=GSE288126).

### Statistical analysis

The results are expressed as the mean of at least three independent measurements, with standard deviations (n=3) presented alongside the average results, unless otherwise stated. All statistical analyses were conducted using GraphPad Prism 8 (GraphPad). A Student’s two-way t-test was employed to make comparisons between different samples. Differences were considered to be statistically significant when *P*≤0.05.

## SUPPLEMENTAL MATERIAL

**Fig. S1 Investigation of involvement of proteasome degradation in E6-induced Top2α expression.** Western blot analysis was used to measure the expression levels of Top2α in LXSN, E6, E7, and E6/E7-overexpressing cells treated with MG132 (10 μM) for 6 hours.

**Fig. S2 Structure and CTD sequence alignment of the CTD of TOP2B and TOP2A.** (A) Structure alignment between TOP2B in green (PDB: 3QX3) and TOP2A in cyan (PDB: 5GWK). (B) Sequence alignment of the CTD of Top2β and Top2α. Red boxes are represented the ubiquitination sites, while blue and green boxes indicate the phosphorylation sites and SUMOylation site, respectively.

**Table. S1.** The protein levels of inflammatory factors in vaginal discharge from normal, CIN and cervical cancer.

|  | Cancer | CIN | Normal |
| --- | --- | --- | --- |
| CCL21 | 396158.6 | 149624.96 | 220264.3 |
| CXCL9 | 19495.74 | 8380.7211 | 3084.577 |
| CXCL6 | 10166.22 | 8907.6017 | 13372.42 |
| IL-6 | 4363.257 | 1312.2528 | 1467.513 |
| CCL1 | 155.54 | 30.692222 | 37.83833 |
| IFN-gamma | 483.3889 | 202.74722 | 161.1661 |
| CXCL12 | 7659.833 | 4172.9606 | 5111.081 |
| CXCL11 | 2164.221 | 565.61778 | 248.2378 |
| CCL7 | 476.595 | 177.01444 | 165.3817 |
| IL-16 | 8857.167 | 3821.6189 | 4224.788 |
| CCL13 | 2286.141 | 2293.4578 | 2416.746 |
| CCL22 | 851.7117 | 264.27056 | 417.7611 |
| CCL24 | 1301.854 | 560.24222 | 1181.166 |
| GM-CSF | 79.73333 | 42.968333 | 38.86167 |
| MIF | 434924.7 | 135554.94 | 136684.8 |
| TNF-alpha | 458.4644 | 200.37944 | 120.3606 |
| CCL23 | 69.22056 | 50.004444 | 58.97167 |
| IL-2 | 23.31667 | 12.467222 | 11.53444 |
| IL-1beta | 2149.173 | 864.21056 | 566.2378 |
| CCL11 | 1045.079 | 489.46667 | 683.9533 |
| CCL25 | 15963.72 | 6601.62 | 6281.565 |
| CXCL10 | 59327.86 | 18540.232 | 14550.96 |
| IL-4 | 74.59778 | 38.248889 | 44.905 |
| CCL2 | 1496.684 | 635.90167 | 973.5189 |
| IL-8 | >6912 ↑ | 6586.1667 | 6180.789 |
| CCL3 | 648.8144 | 487.71944 | 398.7428 |
| IL-10 | 94.52778 | 81.344444 | 67.15444 |
| CCL8 | 1978.569 | 560.30889 | 267.5911 |
| CXCL1 | 33429.72 | 45772.611 | 61117.22 |
| CCL20 | 1280.647 | 496.05444 | 710.925 |
| CXCL16 | 496.9483 | 145.89111 | 316.1372 |
| CCL26 | 1812.824 | 633.73333 | 590.5567 |
| CCL15 | 2670.074 | 1168.0528 | 1861.291 |
| CCL17 | 111.0183 | <1.60 ↓ | <1.60 ↓ |
| CCL27 | 64.90556 | 15.701667 | 20.60056 |
| CXCL5 | 27848.5 | 40111.5 | 29944.61 |
| CXCL13 | 3108.459 | 1012.7406 | 2133.86 |
| CCL19 | 958.0806 | 673.87 | 680.0689 |
| CX3CL1 | 7628.78 | 2011.5178 | 1821.803 |
| CXCL2 | 32842.28 | 24513.949 | 30591.62 |

**Table. S2.**
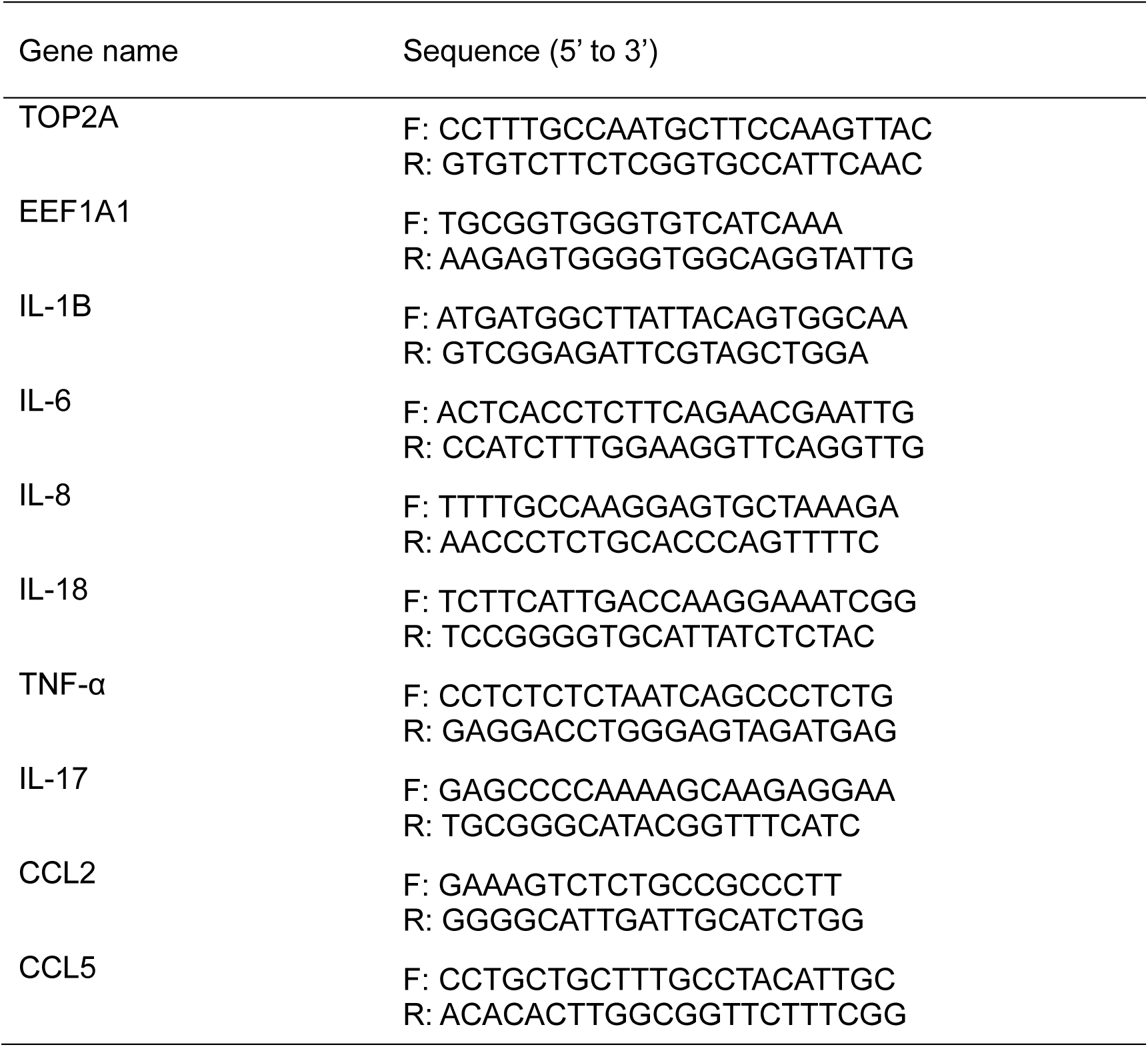
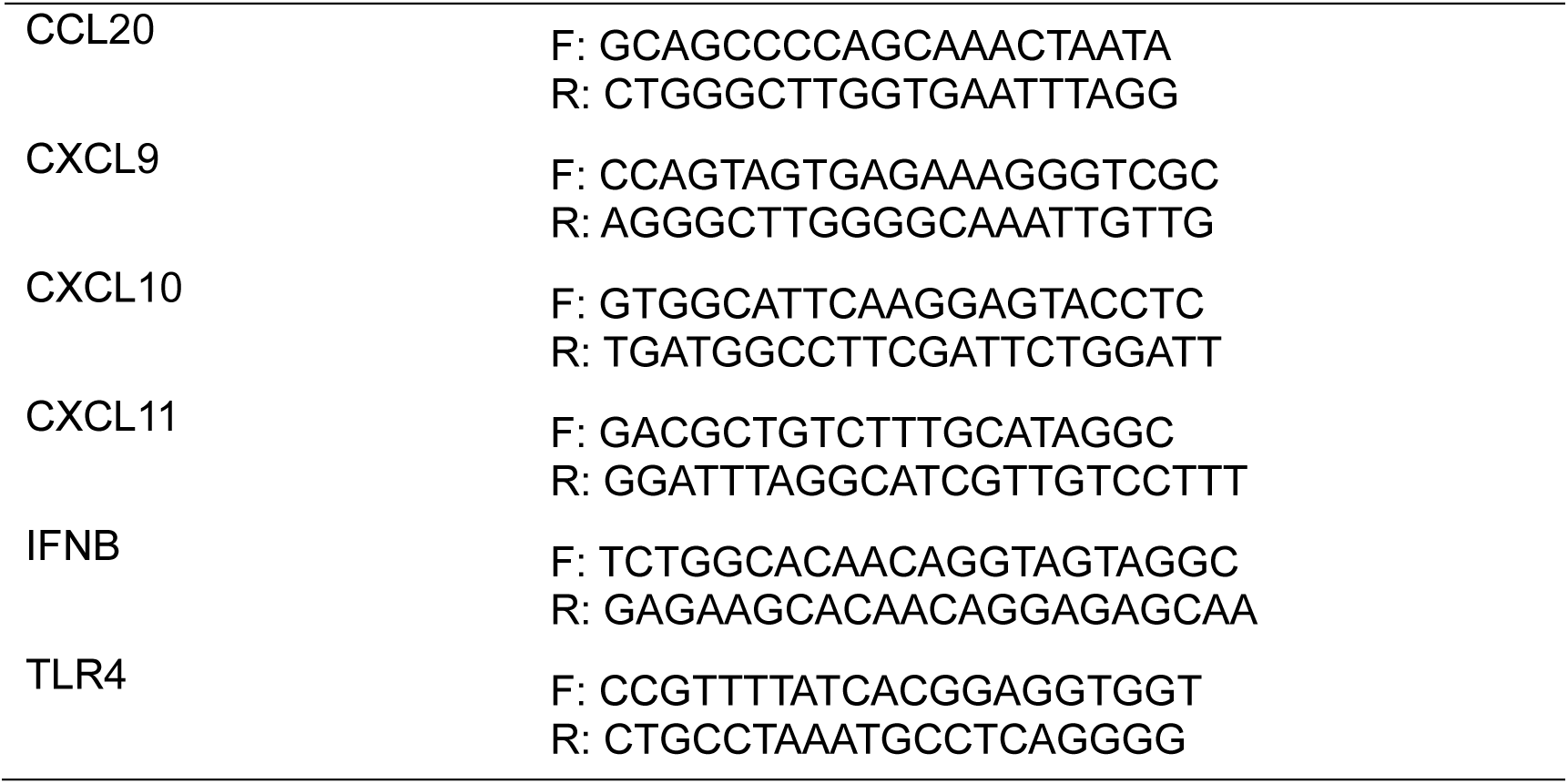
RT-qPCR primer sequences.

**Table. S3.**
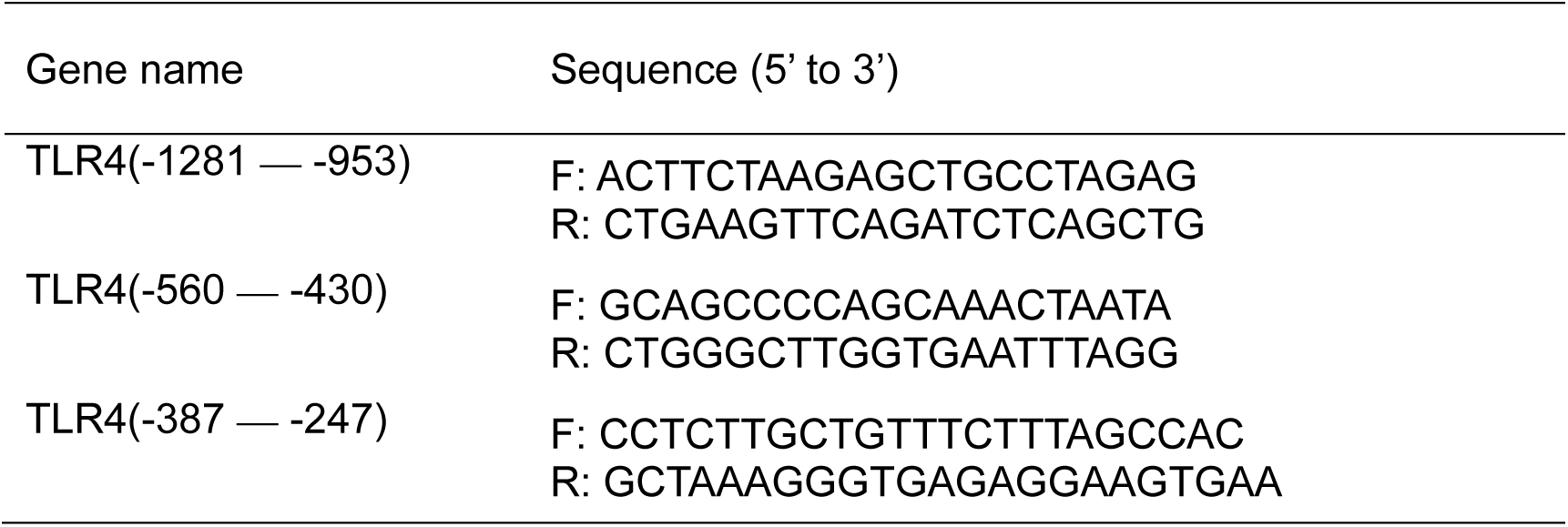
ChIP-qPCR primer sequences.

**Table. S4.**
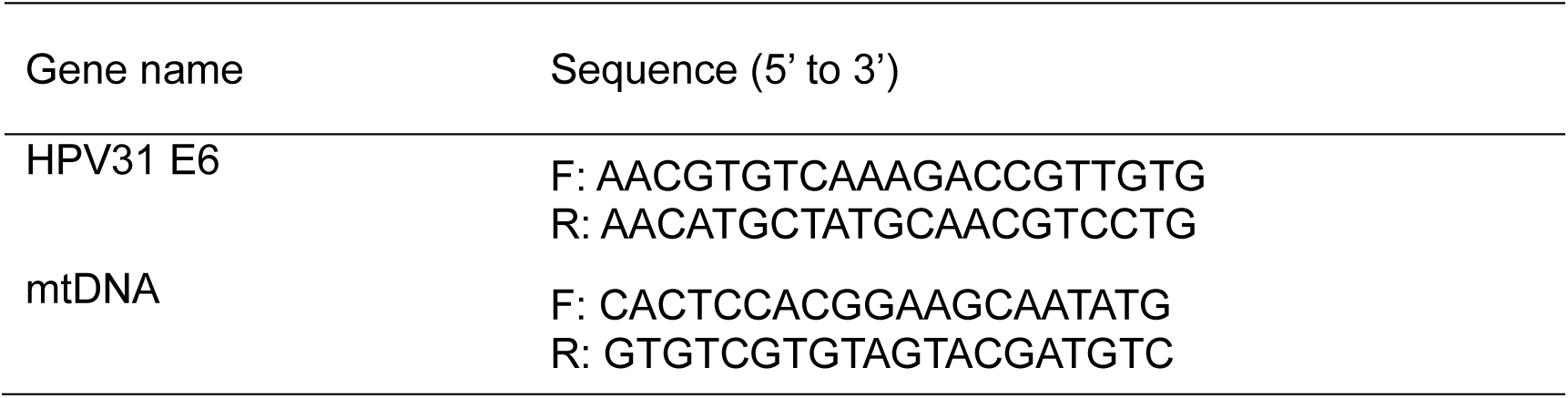
DNA-qPCR primer sequences.

## ETHICS APPROVAL

The clinical samples, including human cervical cancer, cervical intraepithelial neoplasia (CIN) and normal tissue, were collected with the approval of the Women’s Hospital, School of Medicine, Zhejiang University Institutional Review Board (IRB-20220235-R).

## DATA AVAILABILITY

High-throughput sequence data were uploaded to the GEO database. The GEO series number is GSE288126. The supplementary Information, including the cytokine microarray and the protein structure analysis, is provided in the Supplementary Materials accompanying this article. Any additional related information about the supplementary data will be provided upon reasonable request.

## ACKNOWLEDGMENTS

Thanks to all members of Hong’s lab for their input. The plasmids pLXSN-vector, pLXSN-31E6, pLXSN-31E7, pLXSN-31E6E7 were kindly provided by Laimins laboratory (Northwestern University, Chicago, USA) and Stubenrauch laboratory (University of Tübingen, Tübingen, Germany).

This work was supported by CQMU Program for Youth Innovation in Future Medicine (172020320220090 W0072) to S. H., Chongqing Oversea Returnee Entrepreneurship and Innovation Programs (RX20092) to S. H., and Natural Science Program of Chongqing Science and Technology Commission (2024NSCQ-KJFZZDX0041) to S. H..

